# Phenotypic response of yeast metabolic network to availability of proteinogenic amino acids

**DOI:** 10.1101/2022.07.19.500570

**Authors:** Vetle Simensen, Yara Seif, Eivind Almaas

## Abstract

Genome-scale metabolism can best be described as a highly interconnected network of biochemical reactions and metabolites. The flow of metabolites, i.e., flux, throughout these networks can be predicted and analyzed using approaches such as flux balance analysis (FBA). By knowing the network topology and employing only a few simple assumptions, FBA can efficiently predict metabolic functions at the genome scale as well as microbial phenotypes. The network topology is represented in the form of genome-scale metabolic models (GEMs), which provide a direct mapping between network structure and function via the enzyme-coding genes and corresponding metabolic capacity. Recently, the role of protein limitations in shaping metabolic phenotypes have been extensively studied following the reconstruction of enzyme-constrained GEMs. This framework has been shown to significantly improve the accuracy of predicting microbial phenotypes, and it has demonstrated that a global limitation in protein availability can prompt the ubiquitous metabolic strategy of overflow metabolism. Being one of the most abundant and differentially expressed proteome sectors, metabolic proteins constitute a major cellular demand on proteinogenic amino acids. However, little is known about the impact and sensitivity of amino acid availability with regards to genome-scale metabolism. Here, we explore these aspects by extending on the enzyme-constrained GEM framework by also accounting for the usage of amino acids in expressing the metabolic proteome. Including amino acids in an enzyme-constrained GEM of *Saccharomyces cerevisiae*, we demonstrate that the expanded model is capable of accurately reproducing experimental amino acid levels. We further show that the metabolic proteome exerts variable demands on amino acid supplies in a condition-dependent manner, suggesting that *S. cerevisiae* must have evolved to efficiently fine-tune the synthesis of amino acids for expressing its metabolic proteins in response to changes in the external environment. Finally, our results demonstrate how the metabolic network of *S. cerevisiae* is robust towards perturbations of individual amino acids, while simultaneously being highly sensitive when the relative amino acid availability is set to mimic a priori distributions of both yeast and non-yeast origins.

## 1 INTRODUCTION

Constraint-based analysis of genome-scale metabolic models (GEMs) have become an integral framework for predicting metabolic phenotypes of biochemical networks at the genome scale (Bordbar et al., 2014; Fang et al., 2020). Here, the network topology of all metabolic transformations believed to occur within an organism is transformed into a stoichiometric and mathematically structured framework. By formulating mass balance constraints from the reaction stoichiometries and investigating their steady-state behavior, the attainable reaction fluxes span a high-dimensional hyperspace containing the set of feasible flux distributions (Orth et al., 2010). This solution space can be analyzed by various techniques, such as random flux sampling (Wiback et al., 2004), elementary flux modes (Schuster et al., 1999), and flux balance analysis (FBA) related approaches (Orth et al., 2010). A fundamental challenge with this formulation, however, is the considerable size of the solution space and the underlying uncertainty in prediction (Bernstein et al., 2021). Consequently, extensive efforts have been invested into increasing the predictive accuracy of constraint-based analysis of GEMs by formulating and integrating additional biological and physio-chemical constraints on the metabolic network to further reduce the size of this solution space. These constraints have included adjusting the reversibilities of reaction fluxes using principles of thermodynamics (Henry et al., 2006), integration of measured exchange fluxes and growth rates (Sulheim et al., 2020), and employing various omics data to constrain reaction fluxes or construct strain- and condition-specific models (Hyduke et al., 2013).

Enzyme-constrained GEMs have become an important addition to this plethora of computational approaches. Here, the fluxes of biochemical reactions are explicitly limited by the availability and kinetic efficiency of the catalyzing enzymes (Chen and Nielsen, 2021). In other words, weights corresponding to capacity constraints are explicitly added to the network. Many different methods have been developed to account for these limitations in flux magnitudes (Beg et al., 2007; Adadi et al., 2012; Sánchez et al., 2017), where GECKO (*G*enome-scale metabolic models with *E*nzymatic *C*onstraints using *K*inetic and *O*mics data) has become one of the more prominent players in the field (Sánchez et al., 2017; Chen and Nielsen, 2021). In GECKO, the metabolic reactions of the GEM are modified to include pseudo-metabolites representing enzyme usage. These pseudo-metabolites are by themselves constrained through the integration of experimental proteomics measurements or by an overall constraint on the cellular allocation of metabolic proteins. With the latter strategy, the GECKO framework is able not only to predict biochemical fluxes of the metabolic network, but also the necessary enzyme levels required to attain a particular metabolic phenotype (Moreno-Paz et al., 2022). When applying the GECKO approach on a GEM of *Saccharomyces cerevisiae*, the flux variability was considerably reduced compared to that of the purely metabolic GEM, and the enzyme-constrained model showed major improvements in growth rate and secretion profile predictions across a range of conditions (Sánchez et al., 2017; Moreno-Paz et al., 2022). Additionally, the model gained the ability to correctly simulate overflow metabolism (i.e., the Crabtree effect) at higher growth rates without imposing any auxiliary *ad hoc* constraints, substantiating the hypothesis that protein limitation, i.e. network-link capacity constraints, is an underlying cause for this physiological behavior.

Enzymes are biological polymers built from amino acids, and the sequence of amino acids is the key determinant on how proteins fold into a functional three-dimensional structure *in vivo* (Whisstock and Lesk, 2003). These spatial configurations give rise to a vast range of biological functions through their impact on a protein’s ability to bind and interact with other biomolecules (e.g., substrate metabolites in biochemical reactions). The relative amino acid composition of predicted proteomes has been shown to sharply segregate species according to distinct phylogenetic clusters (Tekaia and Yeramian, 2006). Additionally, experiments show that the relative amino acid composition is, for the most part, conserved across environmental conditions for *S. cerevisiae* (Chen and Nielsen, 2022). Being one of the largest and most differentially expressed proteome sectors in *S. cerevisiae* (Yu et al., 2020), metabolic proteins constitute a major sink for proteinogenic amino acids. Here, we explore the role of these amino acids in supporting optimal metabolic phenotypes and investigate how perturbations in their availability affects utilization of the yeast metabolic network at the genome-scale.

Expanding the GECKO framework, we formulate a constraint-based metabolic approach, acidFBA, which explicitly accounts for the absolute usage of proteinogenic amino acids to express a given metabolic proteome. This is done by introducing amino acid drain reactions which concomitantly distribute flux from the growth-limiting protein pool towards individual metabolic proteins at stoichiometric amounts given by their respective amino acid sequences. As a proof of concept, we implement our method using an enzyme-constrained GEM of *S. cerevisiae* and investigate the amino acid usage across a range of growth conditions, demonstrating the amino acid distribution to be contingent on both the metabolic phenotype and nutrient availability. By exploring the near-optimal flux phenotype across a range of conditions, we also discover a significant degree of robustness within the metabolic network when faced with perturbations in the availability of individual amino acids. We further demonstrate how this resilience is directly linked to inhomogeneous distributions of protein-bound amino acids across the metabolic enzymes. Lastly, we find that the growth phenotype of the yeast metabolic network is very sensitive when the amino acid availability is hard-constrained to mimic the experimental amino acid profile of both yeast and other non-yeast organisms.

## 2 MATERIALS AND METHODS

### 2.1 AcidFBA formulation

In the following, we provide a brief summary of the GECKO formalism and how acidFBA is constructed as an expansion of the same methodology with regards to proteinogenic amino acid usage. For a more comprehensive description of GECKO and its underlying assumptions, we refer the reader to Ref. (Sánchez et al., 2017).

The standard FBA problem can be expressed as the following linear program

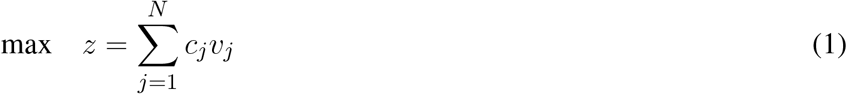

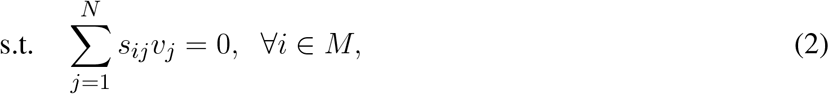

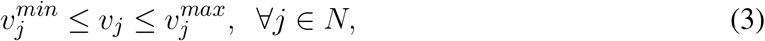

where *v*_*j*_ is the flux (mmol gDW^-1^ h^-1^) through reaction *j, s*_*ij*_ is the stoichiometric coefficient of metabolite *i* in reaction *j, c*_*j*_ is the relative contribution reaction *j* to the objective function, 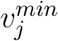 and 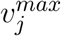 denote the lower and upper bounds on the flux through reaction *j*, while *M* and *N* denote the sets of metabolites and reactions, respectively.

Commonly, 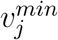 and 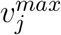 are set at arbitrarily high values to prevent these from being growth limiting. However, the maximal flux (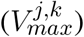) of reaction *j* catalyzed by a given enzyme *k*, as defined by classical Michaelis-Menten-like kinetics in the saturation regime, is determined by the product of enzyme concentration, [*E*_*k*_] (mmol gDW^-1^), and the corresponding catalytic turnover number, 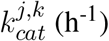

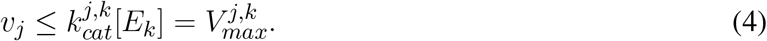

As the cell has a finite amount of available metabolic enzymes, one may formulate a global protein constraint using the following inequality

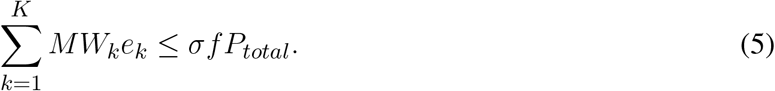

Here, *MW*_*k*_ is the molecular weight (g mol ^-1^) of protein *E*_*k*_, *e*_*k*_ is the flux through the enzyme source reaction (in units mmol gDW^-1^), *P*_*total*_ is the total protein fraction in the cell (g gDW^-1^), *f* is the mass fraction of proteins that are accounted for in the model, while *σ* is the average substrate saturation of enzymes *in vivo*. In GECKO, this is implemented by adding a sink reaction supplying the model with a protein pseudo-metabolite (flux in units g gDW^-1^) representing the aggregated sum of available enzyme mass

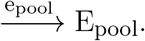

We add *K* enzyme pseudo-reactions to the model, all of which draw from this enzyme pool (flux in units mmol gDW^-1^) towards each corresponding enzyme *E*_*k*_

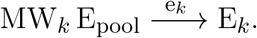

Constraining the flux through the enzyme pool reaction by the total amount of available metabolic proteins, we get the following inequality

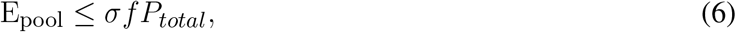

which corresponds directly with that of Eq. (5). Each enzyme *E*_*k*_ is then included as a substrate in the reactions they catalyze, using their reaction-specific inverse turnover number 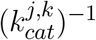 as its stoichiometric coefficients.

In acidFBA, we also consider the necessary usage of proteinogenic amino acids for expressing this metabolic proteome. Each protein is composed of a specific amino acid composition. The flux through the protein sink is therefore allocated into *L* = 20 individual amino acid drain reactions (flux in units g gDW^-1^) of the form

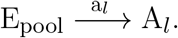

Each enzyme *E*_*k*_ draws from these *L* amino acid drains by introducing the following reactions

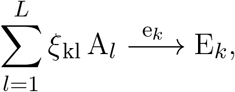

with stoichiometric coefficients given by the elements of the amino acid composition matrix *ξ*. Note that the amino acids *A*_*l*_ are distinct entities from the amino acid metabolites of the GEM. Each entry *ξ*_*kl*_ denotes the molecular weight of amino acid *A*_*l*_ with respect to a given enzyme *E*_*k*_, i.e., gram amino acid per mmol protein (g mmol^-1^), such that the flux *e*_*k*_ has units of mmol protein per gram dry weight (mmol gDW^-1^). Using these sets of reactions, we can formulate the following mass balance for every amino acid *A*_*l*_

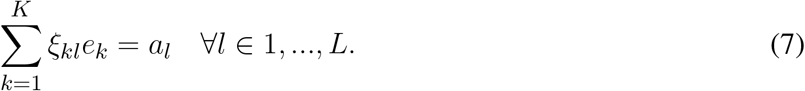

Adapting Eq. (6) to account for these amino acid drains, we obtain an analogous inequality constraint defining the acidFBA framework

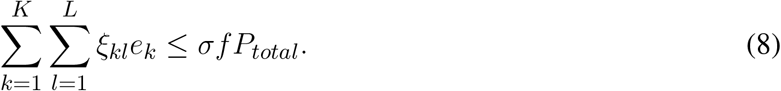

### 2.2 Reconstruction of an acidFBA-GEM

We constructed an acidFBA-GEM of *S. cerevisiae* from the consensus enzyme-constrained GEM (ecGEM) Yeast v. 8.3.4 (Domenzain et al., 2018). Sequence information for all model proteins were retrieved from the UniProt database (Bateman et al., 2021) (UPID UP000002311) and was used to construct the amino acid composition matrix *ξ* using the polymeric molecular formulas for every amino acid. The existing protein pseudo-reactions were replaced by analogous amino acid pseudo-reactions, drawing flux from the enzyme pool reaction towards each metabolic protein at stoichiometric amounts determined by the elements of *ξ*. Finally, an amino acid drain reaction was added for each of the 20 proteinogenic amino acids, bridging the enzyme pool with the metabolic proteins of the model.

### 2.3 Computational modeling of exponential growth and amino acid levels

To simulate early exponential growth in batch culture, we used the protein availability as the only growth-limiting constraint, assuming a total cellular protein fraction *P*_*total*_ = 0.5 g gDW^-1^ (Sánchez et al., 2017). The amino acid levels were calculated by constraining the biomass flux at optimum, then minimizing the overall sum of fluxes. We used absolute quantitative proteomics data from Ref. (Di Bartolomeo et al., 2020) to compare the simulated amino acid profile of the GECKO-implemented proteins. The allowable variability of amino acids was simulated using flux variability analysis (FVA) at 99% of optimal growth to avoid numerical errors. These flux ranges were subsequently normalized by the corresponding mean flux (g gDW^-1^), awarding comparable relative variability scores for each amino acid.

### 2.4 Computational modeling of growth-rate dependency on amino acid usage

The growth-rate dependency of the proteinogenic amino acid distribution was investigated by simulating chemostat growth. Briefly, we constrained the growth rate and limited the availability of metabolic proteins by fitting a non-linear regression model to growth-rate dependent protein mass-fractions of biomass from chemostat cultivations of *S. cerevisiae* (Pejin and Razmovski, 1993). As proposed by Ref. (Sánchez et al., 2017), we further added flux constraints on the exchanges of pyruvate, (R,R)-2,3-butanediol, acetaldehyde, and glycine to physological levels, as well as blocking the intracellular transport of L-serine from the mitochondrial to the cytoplasmic compartment (details can be found in the aminoAcidGrowthRates.m file in the git repository). The enzyme usage was then minimized to obtain a unique distribution of amino acids.

The amino acid distributions of a fully fermentative and a fully respiratory metabolism were simulated in a similar way to that of batch growth, as described previously. In addition to removing the availability of oxygen, anaerobic conditions were achieved by changing the biomass composition, introducing condition-specific constraints, as well as adjusting the growth-associated and non-growth associated maintenance terms (Sánchez et al., 2017). Respiratory conditions were enforced by constraining the specific growth rate to a level where protein availability was non-limiting (arbitrarily set to be 0.2 h^−1^).

### 2.5 Random sampling of nutrient conditions and amino acid sensitivity analysis

To begin with, we identified the set of viable nutrient sources in the model for each of the following elemental classes: carbon, nitrogen, phosphorus, and sulphur. From these viable candidates, we randomly selected *N* = 5, 000 combinations as boundary conditions for the subsequent simulation of optimal growth phenotypes. Limitations on the uptake fluxes of each combination was removed, and growth was optimized by a standard FBA formulation with the same protein limitation as used for simulating exponential batch growth. Subsequently, the overall sum of fluxes was minimized to obtain a singular distribution of amino acids for each sampled condition. The allowable variability of amino acids at an optimality threshold of 99% was calculated by FVA as described previously.

### 2.6 Robustness analysis of alternative amino acid distributions

The amino acid mass distributions of the non-yeast organisms were retrieved from the biomass objective functions of their respective GEMs deposited in the BiGG database (King et al., 2015): iML1515 for *Escherichia coli* (Monk et al., 2017), iYO844 for *Bacillus subtilis* (Oh et al., 2007), and iJN1462 for *Pseudomonas putida* (Nogales et al., 2020). The amino acid distribution of *S. cerevisiae* was acquired from experimental measurements of the GECKO-implemented proteins (Di Bartolomeo et al., 2020). Initially, the relative amino acid distribution of yeast was included as flux constraints on each amino acid drain reaction of the acidFBA-GEM. The maximal growth of the model was identified by performing a standard FBA, providing a reference optimal growth state. Secondly, a hard constraint for the biomass reaction flux was introduced at a fraction of this reference optimal growth rate. Then, for each organism individually, a unique amino acid profile was simulated by minimizing the Euclidean distance between the amino acid profile of the acidFBA-model and the species-specific amino acid distribution. The growth fraction was iteratively incremented within an interval from 0.1 to an upper value defining the relative improvement in growth rate of the acidFBA-model without any constraints on the amino acid profile compared to the reference state. Finally, the absolute relative differences between the species-specific amino acid distributions and the simulated amino acid profiles were calculated.

## 3 RESULTS

### 3.1 AcidFBA accurately predicts metabolic amino acid profiles

We constructed an acidFBA-GEM of *S. cerevisiae* metabolic network by adapting the ecGEM Yeast v. 8.3.4 to account for proteinogenic amino acid requirements for expressing its metabolic proteome. Using absolute quantitative proteomics from Ref. (Di Bartolomeo et al., 2020) of *S. cerevisiae* strain CEN.PK113-7D sampled during exponential growth on glucose, we compared the *in vivo* and *in silico* profile of amino acid usage. Figure 1 shows that the amino acid profile of the acidFBA-GEM displays a strong correlation with that of the experimental measurements of the GECKO-implemented proteins (*R*^2^ = 0.96, *p* = 6.8e^−14^). This correspondence is further substantiated in the predicted enzyme mass employed by the model and that of the experimental data, where the overall protein usage *in vivo* and *in silico* was found to be 0.0928 and 0.0979 g gDW^−1^, respectively. When also accounting for the mass contribution of the metabolically active (i.e., model-predicted at non-zero levels), but unmeasured GECKO proteins (0.0059 g gDW^−1^ in silico), the simulated enzyme mass fraction closely mirror that of the experimental data.

**Figure 1.**
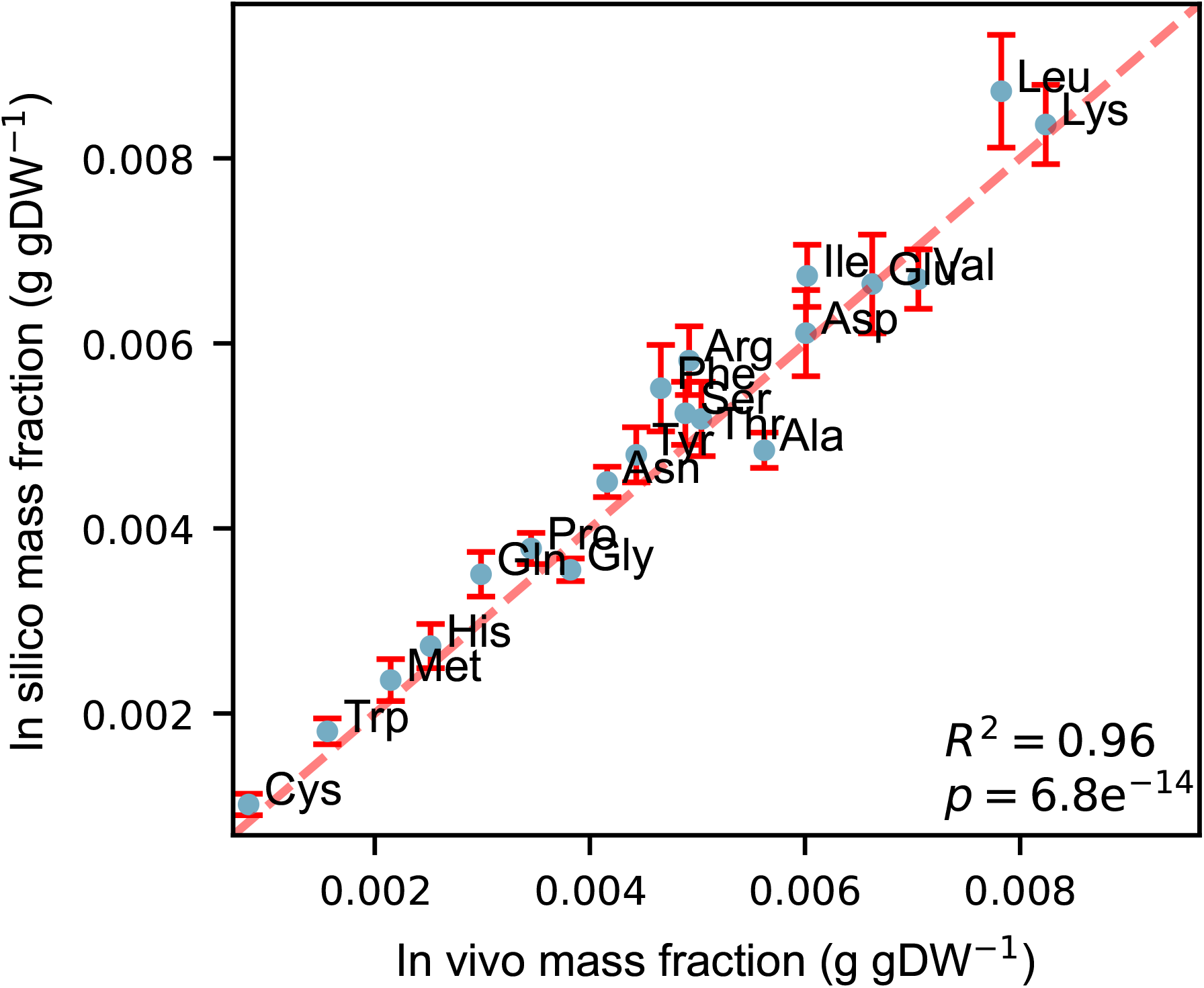
Correlation between *in vivo* and *in silico* amino acid mass fractions (g gDW^-1^) during exponential growth on a defined, minimal glucose medium. Experimental mass fractions were estimated from the GECKO-implemented proteins only. Error bars denote the simulated variability of amino acid usage at 99% of the optimal objective value.

We also simulated the allowable flux variability to quantitatively investigate the flexibility of amino acid supply in supporting a close-to optimal growth phenotype. These flux ranges were simulated by performing FVA on the amino acid drains at 99% of the optimal objective value (error bars in Figure 1). Normalizing the resulting flux ranges by their mean flux values, we found that the relative variability of all amino acids greatly exceeded the selected deviation from growth optimality (S1 fig), displaying a mean relative variability of ∼ 4%. Certain amino acids, such as tryptophan and cysteine, displayed deviations at approximately 8% while effectively maintaining the same growth phenotype through the use of alternative subsets of the metabolic network with a different proteinogenic amino acid distribution (S1 fig).

### 3.2 Amino acid levels are not invariant to changes in metabolic phenotypes

Microbes adjust the expression of their metabolic proteome in response to changes in growth conditions and availability of extracellular nutrients, thus improving chances for survival and proliferation (Schmidt et al., 2016). Through our inclusion of amino acid usage within an enzyme-constrained framework, we are able to explore the variation in proteinogenic amino acid requirements under various nutrient and growth conditions.

To investigate the changes in relative amino acid utilization under varying specific growth rates, we simulated the growth-rate dependency of amino acid usage on a minimal glucose media by constraining the biomass flux (i.e., growth rate) and subsequently minimizing the flux through the protein pool reaction (i.e., overall use of metabolic protein). While akin to parsimonious FBA (pFBA) (Lewis et al., 2010), where the sum of fluxes through all gene-associated reactions are minimized at optimal growth to approximate efficient enzyme usage, this choice of objective allows for a more explicit accounting of catalytic efficiency and proteomic cost by directly minimizing the use of metabolic proteins, i.e. the metabolic network capacity weights.

We find that the relative distribution of amino acids largely remains constant over the growth rates at which the the acidFBA-GEM is predicting a fully respiratory metabolism (Figure 2A, non-shaded area). In this region, we find that the system is unconstrained by the availability of metabolic proteins, and that it employs the most energy efficient metabolic network pathways to convert accessible nutrients to biomass precursors. However, once the growth rate exceeds a critical value of around 0.33 h^−1^, the available enzyme mass become growth-limiting, and the optimal use of the acidFBA-GEM is to reduce the flux through the energy efficient, but protein inefficient, pathways of respiratory metabolism, and concurrently increase the flux through the energy inefficient, but protein efficient, pathways of fermentative metabolism. This transition into the region of proteome-limited growth also entails a shift in the relative distribution of amino acids (Figure 2A). The onset of the prototypical mixed respiro-fermentative metabolism (Crabtree effect) at higher growth rates caused by the concurrent activation and reduction of fermentative and respiratory metabolism, respectively, therefore appears to noticeably impact the relative levels of amino acids needed to express the metabolic proteome. This difference in levels of amino acids is even more apparent when comparing the amino acid distribution of a purely respiratory and a purely fermentative metabolism (Figure 2B). We find that the absolute relative deviation exhibited a median value of around 7%.

**Figure 2.**
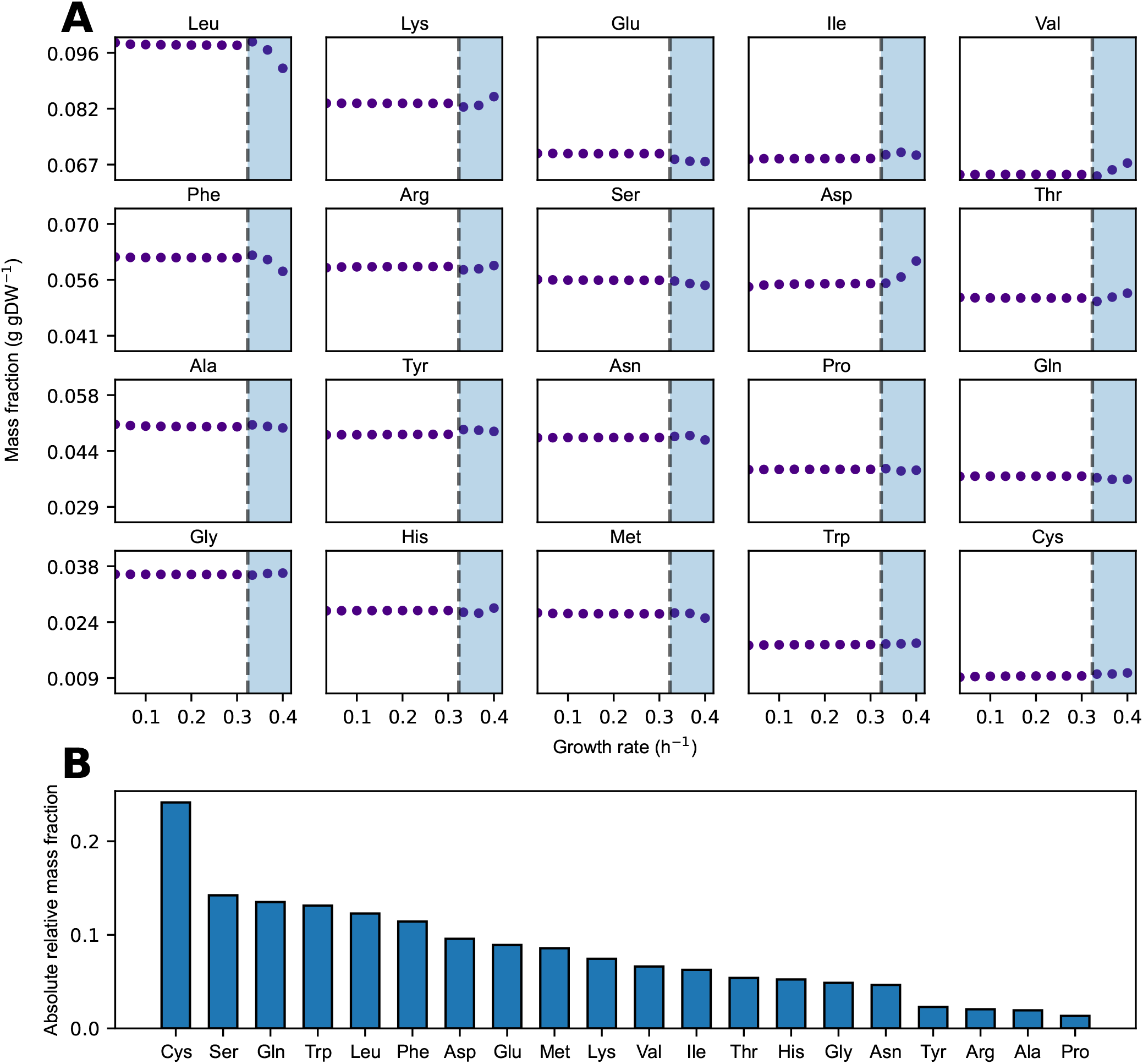
**(A)** Simulated mass fraction (g gDW^-1^) of amino acids in the acidFBA-GEM at varying growth rates. For each simulation, the growth rate was constrained and the resulting flux phenotype was predicted my minimizing the overall enzyme usage. Shaded areas denote the region of protein-limited growth. **(B)** Rank-ordered, absolute relative amino acid mass-fraction deviations of the acidFBA-GEM of a fully fermentative versus a fully respiratory metabolism.

### 3.3 Proteinogenic amino acid usage is contingent on the external nutrient environment

The apparent growth-rate dependency of amino acid usage begs the question to what degree the relative distribution of protein-bound amino acids would need to be modulated in response to condition-specific employment of distinct groups of pathways in the metabolic network. To investigate this, we simulated the amino acid usage on *N* = 5, 000 randomly selected nutrient combinations under protein-limited optimal growth conditions. These *N* = 5, 000 combinations were obtained from a set of 119 carbon sources, 77 nitrogen sources, 47 phosphorus sources, and 14 sulphur sources. While representing only a small fraction of the space of possible combinations, the amino acid distributions were found to converge well to the final distributions presented in Fig. 3. For every combination, the optimal flux phenotype was simulated by constraining the growth rate at the maximal value and subsequently minimizing the overall sum of fluxes. The result of this approach is a unique distribution of amino acids for every simulated condition. We assessed the corresponding variability of the amino acid usage by calculating the relative deviation in mass fractions between every sampled condition versus that of the mean across all samples.

**Figure 3.**
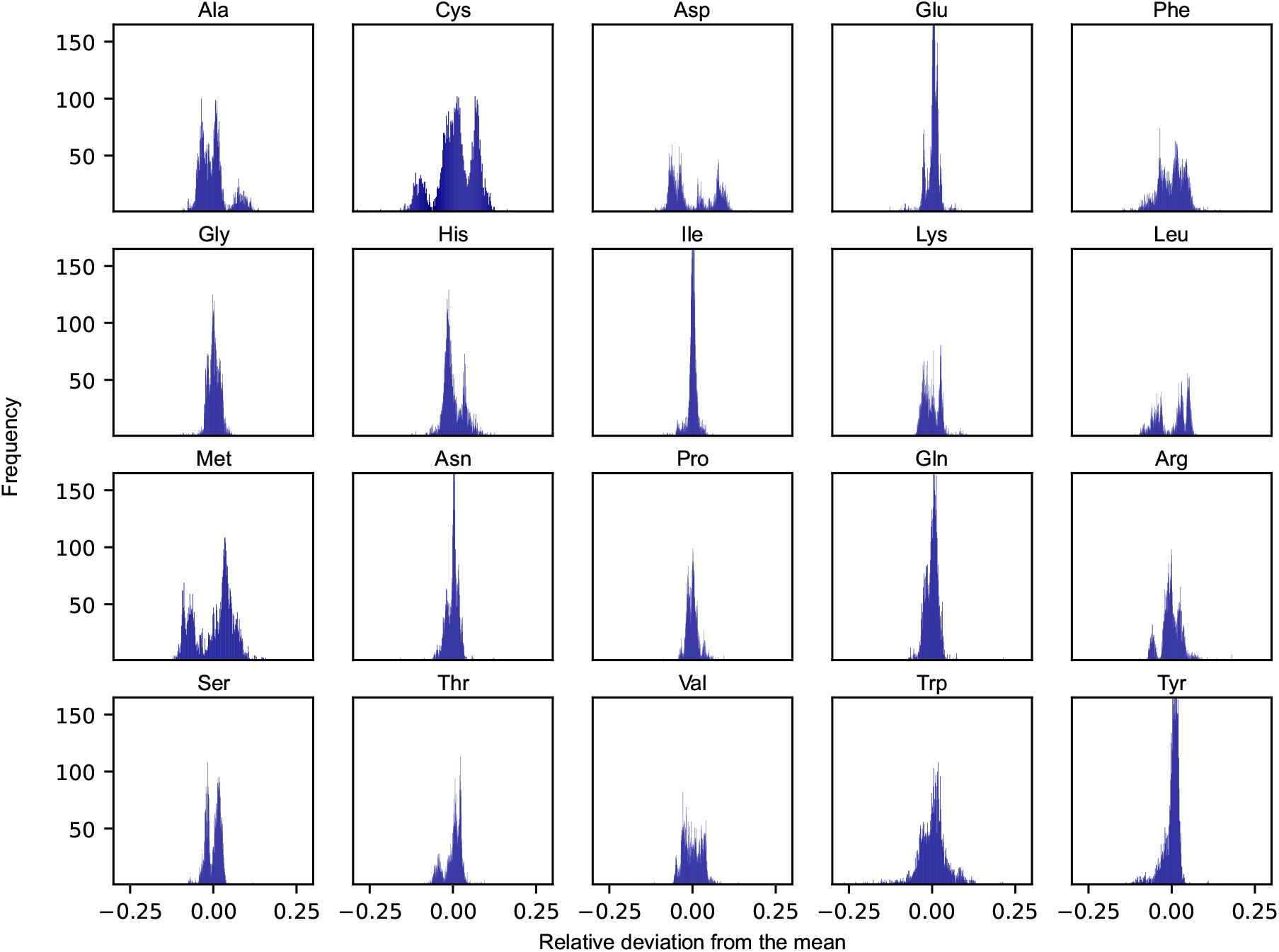
Distributions of the relative deviations from the mean of proteinogenic amino acids across *N* = 5, 000 randomly sampled nutrient conditions. Each condition was defined by a unique combination of viable nutrient sources belonging to each of the four elemental classes: carbon, nitrogen, phosphorus, and sulphur.

As evident from Figure 3, the degree to which a given amino acid is shown to differ across conditions varies substantially. The distributions of some of the individual amino acid are rather narrow across the set of simulated conditions, being largely unaffected by the changing boundary conditions and generally appear to adhere to a species-specific amino acid distribution (e.g., glutamate and isoleucine). Other amino acids, however, are noticeably more variable across the sampled conditions (e.g., cysteine, phenylalanine, and methionine). Interestingly, some of these amino acids also display rather unique and varied distributions, some even in the form of more complex bimodal (e.g., aspartate, leucine, and methionine), or even trimodal distributions, as is the case of cysteine.

### 3.4 Heterogeneous monomer distributions of metabolic proteins are key for robustness against perturbations in amino acid availability

Robustness is a universal property of complex biological systems, and it is fundamental for the maintenance of biological function in the face of internal and external perturbations (Kitano, 2004). The integration of amino acid drains allows us to directly assess the sensitivity of the metabolic phenotype towards perturbations in amino acid availability. These perturbations are directly analogous to evaluating the allowable flux variability of amino acids at close-to optimal growth, as performed on the default minimal glucose media (see Figure 1 and S1 fig).

To form an unbiased view of this metabolic plasticity, we performed FVA on each amino acid drain using the aforementioned *N* = 5, 000 randomly sampled nutrient combinations at a growth optimality threshold of 99%. Mirroring the findings from that of growth on the minimal glucose media, we find that the distribution of feasible mean-normalized flux ranges ubiquitously exceed the chosen deviation from growth optimality for the 20 proteinogenic amino acids. In fact, the median relative variability was found to be ∼ 4.8%, demonstrating an innate buffering capacity of the enzyme-constrained metabolic network on the optimal growth phenotype when subjected to perturbations in the availability of any single amino acid.

The amino acids displaying the greatest flexibility over the simulated nutrient conditions was found to be cysteine (8.6%), methionine (8.1%), glutamate (7.4%), and histidine (6.6%), with the denoted median relative variabilities. Save for glutamate, the relative flexibility of these amino acids also showed a high degree of variation between nutrient conditions (mean standard deviation of 2.7%). Although to a lesser extent, we also found this to be the case for the remaining amino acids, indicating that the magnitude of metabolic robustness is directly contingent on the growth condition.

Next, we sought to explore the underlying causes of this phenotypic robustness by repeating the same condition-dependent sensitivity analysis, but now instead assuming an invariant amino acid distribution for all GECKO-implemented proteins. Using the mean amino acid levels from Figure 1 as a representative distribution, the amino acid composition matrix *ξ* was recalculated and employed to construct a secondary acidFBA-GEM. As evident from the S2 fig, we find that the relative flexibility of each proteinogenic amino acid now directly aligns with the selected deviation from growth optimality (1%). The buffering capacity is thereby lost, as any differential employment of alternative pathways and isozymes has no effect on the relative usage of amino acids. This demonstrates that heterogeneous distributions of protein-bound amino acids in metabolic enzymes is key for this robustness in growth phenotype when faced with perturbations in amino acid availabilities.

### 3.5 Yeast metabolic network is sensitive to a priori constraints on amino acid levels

The relative amino acid profiles of expressed proteomes have been shown to reflect species-specific distributions (Tekaia and Yeramian, 2006). We therefore sought out to evaluate how robust the growth performance of the yeast metabolic network is when the availability of amino acids in the acidFBA framework is set to mimic other unrelated organisms. This analysis was performed on three separate bacterial species, each with a distinct amino acid distribution: *E. coli, B. subtilis*, and *P. putida*. To provide a reference optimal growth state, we enforced the experimental amino acid distribution of yeast (Fig. 1) on the acidFBA-GEM and ran a standard FBA to identify the maximal growth rate. We found this to be approximately 0.15 h^−1^, representing a considerable growth reduction as compared to the default acidFBA-GEM which has no constraints on the relative amino acid profile (maximal growth rate of around 0.38 h^−1^). Interestingly, we found the solution to be infeasible when mimicking the amino acid distributions of the non-yeast species, demonstrating the models inability to grow when the amino acid distribution is hard-constrained and sufficiently dissimilar from the one predicted by the default acidFBA-GEM.

In order to address this, we instead elected to simulate amino acid profiles that were minimally different to the species-specific distributions across a range of relative growth rates (relative to the aforementioned reference optimal growth state). These simulations were performed for all three non-yeast organisms, calculating the absolute relative difference between the amino acid levels of the model and those of the species-specific distributions (Fig. 5). As expected, we found that the non-yeast amino acid distributions are unattainable across all the evaluated growth rates. In order to grow at all, the model needs to considerably adjust its amino acid levels to express the necessary metabolic proteins. For the majority of amino acids, this absolute relative difference is demonstrated to increase in magnitude as a function of the relative growth rate as the amino acid profile of the model approaches the one selected naturally by the model when growing maximally without any amino acid constraints. The same analysis was performed on yeast (blue points in Fig. 5). Here, we observe that the model is able to maintain the experimental amino acid distribution at relative growth rates below 1.0. At this critical point, any incremental increase in growth rate forces the model to adjust its amino acid profile away from this experimental distribution. Unlike the non-yeast species, however, the total redistribution of amino acid levels are generally quite minor.

**Figure 4.**
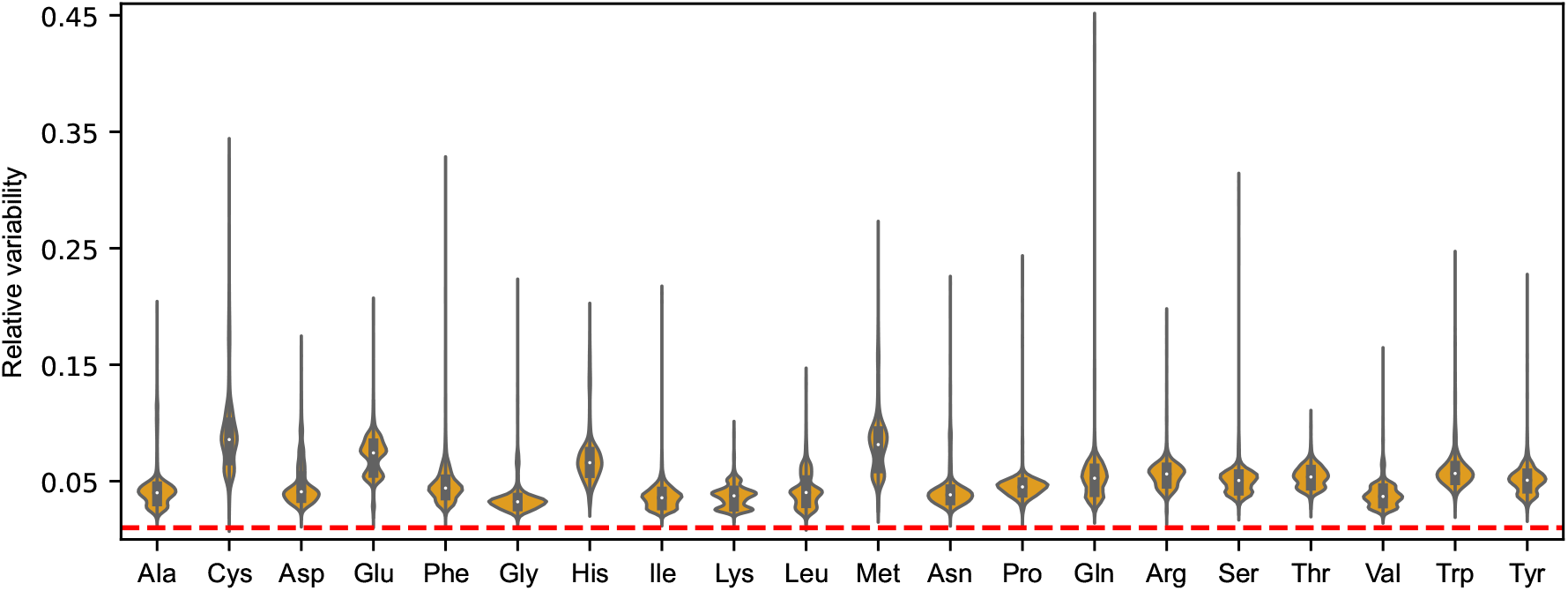
Violin plots of the distributions of mean-normalized flux ranges across *N* = 5, 000 sampled nutrient combinations for the 20 proteinogenic amino acids of the acidFBA model. The feasible flux ranges were simulated by performing a flux variability analysis (FVA) using an optimality threshold of 99%. Dotted line in red denote the selected deviation from growth optimality.

**Figure 5.**
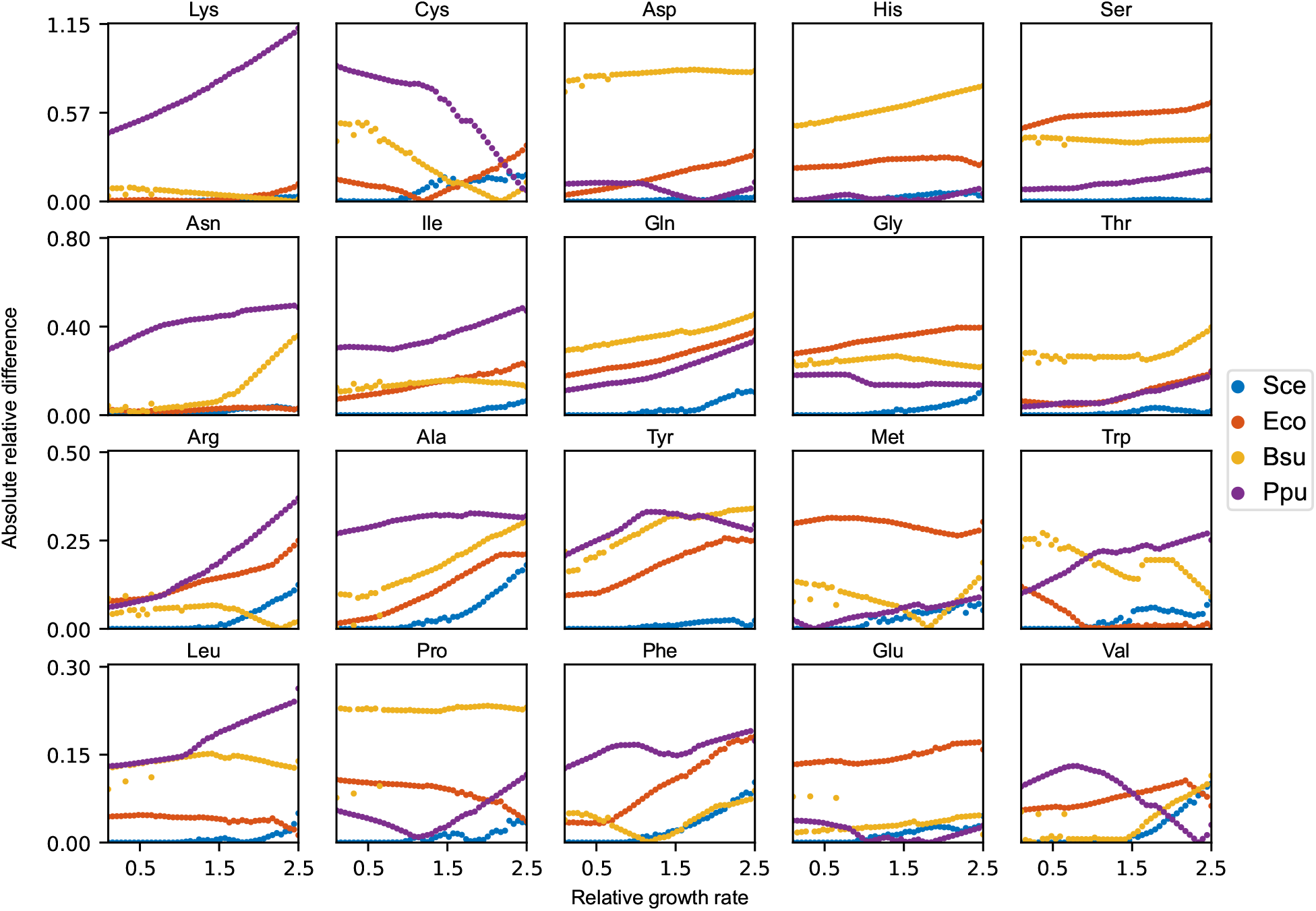
Absolute relative difference of each amino acid between the species-specific amino acid distributions and the simulated amino acid profiles of the acidFBA-GEM at increasing relative growth rates. The simulated profiles were obtained by minimizing the Euclidean distance to the species-specific distributions. Sce: *S. cerevisiae*, Eco: *E. coli*, Bsu: *B. subtilis*, Ppu: *P. putida*.

## 4 DISCUSSION

The integration of protein constraints with GEMs has provided invaluable insight into the role of the cellular proteome in shaping the landscape of feasible metabolic phenotypes (Chen and Nielsen, 2021). By being one of the largest cellular proteome sectors (Liebermeister et al., 2014), metabolic proteins appropriate a considerable proportion of the cellular supply of proteinogenic amino acids. In this work, we extend the protein-limited GEM framework by performing a systematic and quantitative analysis of the role of proteinogenic amino acids in expressing metabolic proteomes to support metabolic phenotypes, and thus, the use and flexibility of metabolic networks. This was implemented by expanding the GECKO formalism by introducing amino acid drains which reroute flux from the growth-limiting protein pool towards the metabolic enzymes of the model. This allows for a direct prediction of the distribution of amino acid mass fractions necessary for achieving a particular metabolic phenotype, as well as assessing the sensitivity of the metabolic network towards perturbations in the availability of amino acids.

While the global amino acid profile of *S. cerevisiae* experimentally has been found to be largely conserved across growth conditions (Chen and Nielsen, 2022), we here show that the metabolically active proteome, as an isolated proteome sector, appears not to exhibit the same degree of invariance when subjected to diverse nutrient environments and growth conditions. In fact, we found that the growth-rate dependency of pathway usage in energy metabolism also impacts amino acid usage, as the fermentative and respiratory pathways are shown to exert different demands on the relative distribution of proteinogenic amino acids (see Figure 2). Moreover, we demonstrate a clear contingency of amino acid levels on the nutrient environment by conducting a large-scale evaluation of amino acid utilization over a wide range of boundary conditions. Here, the relative distribution of the majority of amino acids is shown to display considerable variability across conditions, some even forming characteristic multi-modal distributions. This indicates that a differential expression of metabolic proteins and pathways also entail a corresponding shift and adjustment in the relative levels of necessary proteinogenic amino acids. These findings imply that the *S. cerevisiae* metabolic network must have evolved to efficiently fine-tune its synthesis of proteinogenic amino acids needed for expressing its metabolic proteome in response to fluctuations in the external environment.

By incorporating amino acid levels as variables within the optimization framework, we were able to directly assess the robustness of the metabolic network towards perturbations in the availability of individual amino acids. Interestingly, we find that the metabolic proteome of *S. cerevisiae* has a surprising potential to robustly adapt to perturbations in amino acid availability, as is evident from the feasible flux ranges all exceeding the selected deviation from growth optimality (Figure 4). The effect on the (close-to) optimal growth phenotype by perturbing the supply of any single amino acid is shown to be largely buffered by a redistribution of the remaining amino acids. These reorganizations are caused by compensatory flux rerouting through the use of alternative pathways and isozymes with an altered demand on the particular amino acid. We further showcase how this innate robustness is linked to sequence heterogeneity, and that this ability is altogether lost when the amino acid distributions are assumed universal and invariant across the metabolic proteins of the model. The same tendency for robustness was not observed when the amino acid profile was set to mimic that of unrelated species. In fact, our results suggests that the yeast metabolic network is highly sensitive to any a priori constraints on the relative availability of amino acids. While our results are too insufficient to make grand assertions of broad-scale evolutionary selection pressures for protein sequence heterogeneity, they indicate that cellular implementation of non-homogeneous amino acid distributions has an additional benefit of metabolic robustness when faced with perturbations in amino acid availability.

Yeasts are commonly used as cell factories for the production of heterologous proteins, in particular the synthesis of pharmaceutical proteins that require eukaryal systems for proper folding and post-translational modifications (Baghban et al., 2019). This recombinant protein production subjects the cells to a considerable metabolic burden which consumes important cellular resources necessary for cell growth and maintenance such as metabolic precursors, reducing power, and cellular machineries necessary for all levels of gene expression (Kastberg et al., 2022). The cellular supply of amino acids has been found to be a key limiting factor for translational rates (Gonzalez et al., 2003). Due to the potential differences in amino acid composition of native yeast proteins and the heterologous protein, this expression can potentially affect the availability of specific amino acids (Kastberg et al., 2022). By employing our presented framework, we can thereby explicitly investigate the metabolic effects of expressing any protein with a known amino acid profile and generate well-informed strategies for alleviating these adverse effects on the metabolic network.

As each protein imposes a cost to the organism not only due to peptide bond formation, but also due to the biosynthetic expense of synthesizing its constituent amino acids, the acidFBA formulation also allows for a direct calculation of cellular cost associated with expressing a particular metabolic proteome. It has been demonstrated that highly expressed proteins evolve to utilize less costly amino acids. This was exemplified by Akashi and Gojobori who demonstrated a negative correlation between gene expression and average protein cost when employing codon usage bias as a predictor of translational rates in *Escherichia coli* and *Bacillus subtilis* (Akashi and Gojobori, 2002). Subsequent work in the field added to this hypothesis and showcased the universality of the principle of cost-minimization as a key driver of microbial proteome evolution (Raiford et al., 2008; Seligmann, 2003; Wagner, 2005; Heizer et al., 2006). Recently, Chen and Nielsen demonstrated how the expressed proteome of *S. cerevisiae* tends to minimize the use of amino acids that are proteomically costly to synthesize, rather than those that are costly energetically (Chen and Nielsen, 2022). Consequently, we are of the opinion that AcidFBA is a highly suitable tool for investigating the role of proteomic cost minimization in shaping the metabolic phenotypes, and thereby patterns of metabolic network utilization, of an organism. Additionally, the computational analysis of cost-optimized metabolic states could prove quite valuable for understanding pathway selection in metabolic engineering applications.

## AUTHOR CONTRIBUTIONS

YS and EA conceptualized the study. VS constructed the model and performed all computational analysis under the supervision of EA. VS wrote the first draft of the manuscript. All authors contributed to manuscript revision, read, and approved the submitted version.

## ACKNOWLEDGMENTS

The authors would like to thank Christian Schulz, Emil Karlsen, and Jakob P. Pettersen for helpful discussions and feedback throughout the project.

## DATA AVAILABILITY STATEMENT

All data and necessary code to reproduce the results for this study can be found online in the following Github repository: https://github.com/AlmaasLab/acidFBA.

## REFERENCES

Adadi, R., Volkmer, B., Milo, R., Heinemann, M., and Shlomi, T. (2012). Prediction of Microbial Growth Rate versus Biomass Yield by a Metabolic Network with Kinetic Parameters. PLOS Computational Biology 8, e1002575. doi:10.1371/JOURNAL.PCBI.1002575

Akashi, H. and Gojobori, T. (2002). Metabolic efficiency and amino acid composition in the proteomes of Escherichia coli and Bacillus subtilis. Proceedings of the National Academy of Sciences 99, 3695–3700. doi:10.1073/PNAS.062526999

Baghban, R., Farajnia, S., Rajabibazl, M., Ghasemi, Y., Mafi, A. A., Hoseinpoor, R., et al. (2019). Yeast Expression Systems: Overview and Recent Advances. Molecular Biotechnology 2019 61:5 61, 365–384. doi:10.1007/S12033-019-00164-8

Bateman, A., Martin, M. J., Orchard, S., Magrane, M., Agivetova, R., Ahmad, S., et al. (2021). UniProt: the universal protein knowledgebase in 2021. Nucleic Acids Research 49, D480–D489. doi:10.1093/NAR/GKAA1100

Beg, Q. K., Vazquez, A., Ernst, J., De Menezes, M. A., Bar-Joseph, Z., Barabási, A. L., et al. (2007). Intracellular crowding defines the mode and sequence of substrate uptake by Escherichia coli and constrains its metabolic activity. Proceedings of the National Academy of Sciences of the United States of America 104, 12663. doi:10.1073/PNAS.0609845104

Bernstein, D. B., Sulheim, S., Almaas, E., and Segrè, D. (2021). Addressing uncertainty in genome-scale metabolic model reconstruction and analysis. Genome Biology 2021 22:1 22, 1–22. doi:10.1186/S13059-021-02289-Z

Bordbar, A., Monk, J. M., King, Z. A., and Palsson, B. O. (2014). Constraint-based models predict metabolic and associated cellular functions. Nature Reviews Genetics 2014 15:2 15, 107–120. doi:10.1038/nrg3643

Chen, Y. and Nielsen, J. (2021). Mathematical modeling of proteome constraints within metabolism. Current Opinion in Systems Biology 25, 50–56. doi:10.1016/J.COISB.2021.03.003

Chen, Y. and Nielsen, J. (2022). Yeast has evolved to minimize protein resource cost for synthesizing amino acids. Proceedings of the National Academy of Sciences 119, e2114622119. doi:10.1073/PNAS.2114622119

Di Bartolomeo, F., Malina, C., Campbell, K., Mormino, M., Fuchs, J., Vorontsov, E., et al. (2020). Absolute yeast mitochondrial proteome quantification reveals trade-off between biosynthesis and energy generation during diauxic shift. Proceedings of the National Academy of Sciences of the United States of America 117, 7524–7535. doi:10.1073/PNAS.1918216117/-/DCSUPPLEMENTAL

[Dataset] Domenzain, I., Anton, M., and Sánchez, B. (2018). GitHub - SysBioChalmers/ecModels: A container for all enzyme constrained models created by GECKO.

Fang, X., Lloyd, C. J., and Palsson, B. O. (2020). Reconstructing organisms in silico: genome-scale models and their emerging applications. Nature Reviews Microbiology 2020 18:12 18, 731–743. doi:10.1038/s41579-020-00440-4

Gonzalez, R., Andrews, B. A., Molitor, J., and Asenjo, J. A. (2003). Metabolic analysis of the synthesis of high levels of intracellular human SOD in Saccharomyces cerevisiae rhSOD 2060 411 SGA122. Biotechnology and Bioengineering 82, 152–169. doi:10.1002/BIT.10556

Heizer, E. M., Raiford, D. W., Raymer, M. L., Doom, T. E., Miller, R. V., and Krane, D. E. (2006). Amino Acid Cost and Codon-Usage Biases in 6 Prokaryotic Genomes: A Whole-Genome Analysis. Molecular Biology and Evolution 23, 1670–1680. doi:10.1093/MOLBEV/MSL029

Henry, C. S., Jankowski, M. D., Broadbelt, L. J., and Hatzimanikatis, V. (2006). Genome-Scale Thermodynamic Analysis of Escherichia coli Metabolism. Biophysical Journal 90, 1453. doi:10.1529/BIOPHYSJ.105.071720

Hyduke, D. R., Lewis, N. E., and Palsson, B. O. (2013). Analysis of omics data with genome-scale models of metabolism. Molecular bioSystems 9, 167. doi:10.1039/C2MB25453K

Kastberg, L. L. B., Ard, R., Jensen, M. K., and Workman, C. T. (2022). Burden Imposed by Heterologous Protein Production in Two Major Industrial Yeast Cell Factories: Identifying Sources and Mitigation Strategies. Frontiers in Fungal Biology 0, 1. doi:10.3389/FFUNB.2022.827704

King, Z. A., Lu, J., Dräger, A., Miller, P., Federowicz, S., Lerman, J. A., et al. (2015). BiGG Models: A platform for integrating, standardizing and sharing genome-scale models. Nucleic Acids Research 44, D515–D522. doi:10.1093/nar/gkv1049

Kitano, H. (2004). Biological robustness. Nature Reviews Genetics 2004 5:11 5, 826–837. doi:10.1038/nrg1471

Lewis, N. E., Hixson, K. K., Conrad, T. M., Lerman, J. A., Charusanti, P., Polpitiya, A. D., et al. (2010). Omic data from evolved E. coli are consistent with computed optimal growth from genome-scale models. Molecular Systems Biology 6, 390. doi:10.1038/MSB.2010.47

Liebermeister, W., Noor, E., Flamholz, A., Davidi, D., Bernhardt, J., and Milo, R. (2014). Visual account of protein investment in cellular functions. Proceedings of the National Academy of Sciences of the United States of America 111, 8488–8493. doi:10.1073/PNAS.1314810111/ASSET/412674C7-2411-4D67-A176-C61B53B65021/ASSETS/GRAPHIC/PNAS.1314810111FIG04.JPEG

Monk, J. M., Lloyd, C. J., Brunk, E., Mih, N., Sastry, A., King, Z., et al. (2017). iML1515, a knowledgebase that computes Escherichia coli traits. Nature Biotechnology 35, 904–908. doi:10.1038/nbt.3956

Moreno-Paz, S., Schmitz, J., Martins dos Santos, V. A., and Suarez-Diez, M. (2022). Enzyme-constrained models predict the dynamics of Saccharomyces cerevisiae growth in continuous, batch and fed-batch bioreactors. Microbial Biotechnology 15, 1434–1445. doi:10.1111/1751-7915.13995

Nogales, J., Mueller, J., Gudmundsson, S., Canalejo, F. J., Duque, E., Monk, J., et al. (2020). High-quality genome-scale metabolic modelling of Pseudomonas putida highlights its broad metabolic capabilities. Environmental Microbiology 22, 255–269. doi:10.1111/1462-2920.14843

Oh, Y. K., Palsson, B. O., Park, S. M., Schilling, C. H., and Mahadevan, R. (2007). Genome-scale reconstruction of metabolic network in Bacillus subtilis based on high-throughput phenotyping and gene essentiality data. Journal of Biological Chemistry 282, 28791–28799. doi:10.1074/JBC.M703759200/ATTACHMENT/16CDEC70-9051-4225-A19B-1667C848C556/MMC1.ZIP

Orth, J. D., Thiele, I., and Palsson, B. O. (2010). What is flux balance analysis? Nature biotechnology 28, 245–248. doi:10.1038/nbt.1614

Pejin, D. and Razmovski, R. (1993). Continuous cultivation of the yeastSaccharomyces cerevisiae at different dilution rates and glucose concentrations in nutrient media. Folia Microbiologica 1993 38:2 38, 141–146. doi:10.1007/BF02891696

Raiford, D. W., Heizer, E. M., Miller, R. V., Akashi, H., Raymer, M. L., and Krane, D. E. (2008). Do Amino Acid Biosynthetic Costs Constrain Protein Evolution in Saccharomyces cerevisiae? Journal of Molecular Evolution 2008 67:6 67, 621–630. doi:10.1007/S00239-008-9162-9

Sánchez, B. J., Zhang, C., Nilsson, A., Lahtvee, P.-J., Kerkhoven, E. J., and Nielsen, J. (2017). Improving the phenotype predictions of a yeast genome-scale metabolic model by incorporating enzymatic constraints. Molecular Systems Biology 13, 935. doi:10.15252/MSB.20167411

Schmidt, A., Kochanowski, K., Vedelaar, S., Ahrné, E., Volkmer, B., Callipo, L., et al. (2016). The quantitative and condition-dependent Escherichia coli proteome. Nature Biotechnology 2015 34:1 34, 104–110. doi:10.1038/nbt.3418

Schuster, S., Dandekar, T., and Fell, D. A. (1999). Detection of elementary flux modes in biochemical networks: a promising tool for pathway analysis and metabolic engineering. Trends in Biotechnology 17, 53–60. doi:10.1016/S0167-7799(98)01290-6

Seligmann, H. (2003). Cost-Minimization of Amino Acid Usage. Journal of Molecular Evolution 2003 56:2 56, 151–161. doi:10.1007/S00239-002-2388-Z

Sulheim, S., Kumelj, T., van Dissel, D., Salehzadeh-Yazdi, A., Du, C., van Wezel, G. P., et al. (2020). Enzyme-Constrained Models and Omics Analysis of Streptomyces coelicolor Reveal Metabolic Changes that Enhance Heterologous Production. iScience 23, 101525. doi:10.1016/J.ISCI.2020.101525

Tekaia, F. and Yeramian, E. (2006). Evolution of proteomes: Fundamental signatures and global trends in amino acid compositions. BMC Genomics 7, 1–11. doi:10.1186/1471-2164-7-307/FIGURES/4

Wagner, A. (2005). Energy Constraints on the Evolution of Gene Expression. Molecular Biology and Evolution 22, 1365–1374. doi:10.1093/MOLBEV/MSI126

Whisstock, J. C. and Lesk, A. M. (2003). Prediction of protein function from protein sequence and structure. Quarterly Reviews of Biophysics 36, 307–340. doi:10.1017/S0033583503003901

Wiback, S. J., Famili, I., Greenberg, H. J., and Palsson, B. (2004). Monte Carlo sampling can be used to determine the size and shape of the steady-state flux space. Journal of Theoretical Biology 228, 437–447. doi:10.1016/J.JTBI.2004.02.006

Yu, R., Campbell, K., Pereira, R., Björkeroth, J., Qi, Q., Vorontsov, E., et al. (2020). Nitrogen limitation reveals large reserves in metabolic and translational capacities of yeast. Nature Communications 2020 11:1 11, 1–12. doi:10.1038/s41467-020-15749-0

