## Supplementary Figures for "Phenotypic response of yeast metabolic network to availability of proteinogenic amino acids"

### *Supplementary Material*

#### 1 SUPPLEMENTARY TABLES AND FIGURES

##### 1.1 Figures

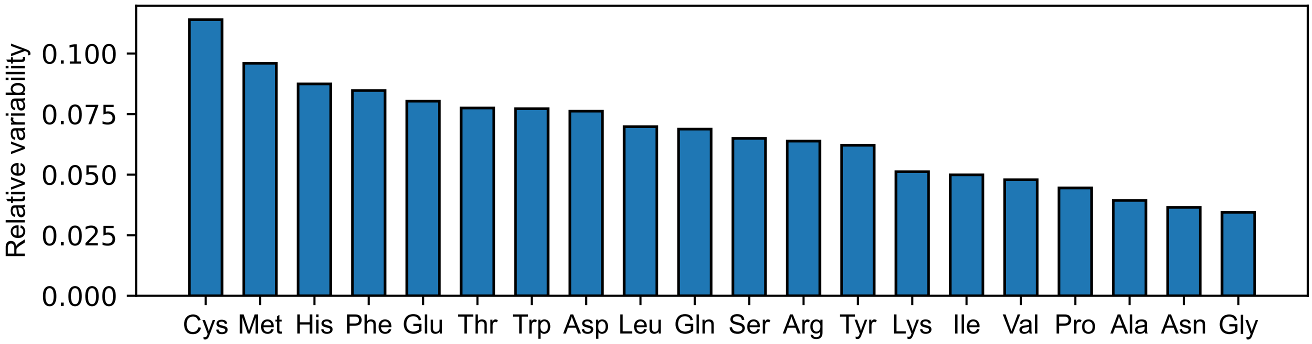


**Figure S1.** Rank-ordered, relative flux variability of proteinogenic amino acids on a defined, minimal glucose medium. Calculated by normalizing the feasible flux ranges at $99\%$ of optimal growth by their corresponding mean flux value. Mean-flux normalized counterpart to the error bars in Fig. 1.

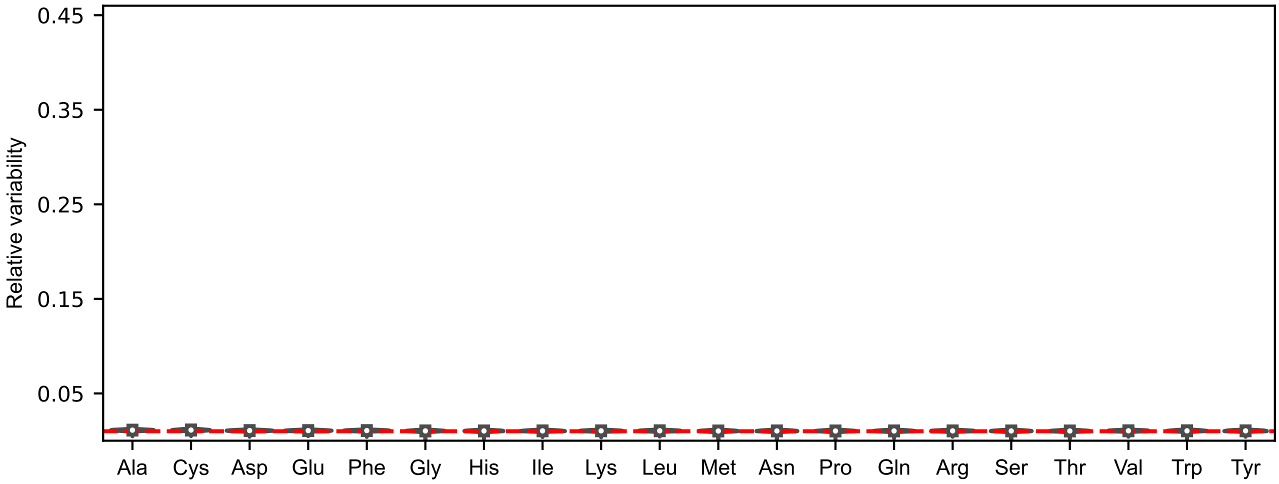


**Figure S2.** Violin plots of the distributions of mean-normalized flux ranges across $N = 5000$ sampled nutrient combinations for the 20 proteinogenic amino acids of the acidFBA-GEM using an invariant amino acid distribution for all GECKO-implemented proteins. The feasible flux ranges were simulated by performing a flux variability analysis (FVA) using an optimality threshold of $99\%$. Dotted line in red denote the selected deviation from growth optimality.
